# Hypoxia and Associated Acidosis Generate Cell-Type Specific Myeloid Responses in Glioblastoma

**DOI:** 10.64898/2026.02.26.707379

**Authors:** Aliisa M. Tiihonen, Iida Salonen, Iina Koivisto, Anni S. Ritamäki, Serafiina Jaatinen, Tanja Hyvärinen, Johanna Tilvis, Joose M. Kreutzer, Göktug Karabiyik, Sonja Mäntylä, Maryam Mohammadlou, Miina Hoikka, Masi Valkonen, Jürgen Beck, Roland Rölz, Mikael Marttinen, Joonas Haapasalo, Matti Nykter, Pekka Ruusuvuori, Seppo Parkkila, Pasi Kallio, Sanna Hagman, Juha Kesseli, Vidhya M. Ravi, Hannu Haapasalo, Arja Jukkola, Kevin Joseph, Kirsi J. Rautajoki

## Abstract

Hypoxia is a defining feature of glioblastoma (GBM), yet how it cooperates with hypoxia-associated acidosis to shape microglia and infiltrating monocyte-derived macrophages (MDM) remains poorly understood. We integrated cyclic immunohistochemistry, single-cell RNA sequencing, spatial transcriptomics, *in vitro* cell cultures, and DNA methylation profiling to outline hypoxia-driven responses in up to 136 GBMs. These hypoxic niches were selectively enriched for MDMs that activated carbonic anhydrase (CA) mediated pH buffering and other metabolic adaptation programs, enabling survival in acidic hypoxia, increasingly interacted with cancer cells, and show polarization toward immunosuppressive myeloid-derived suppressor cell (MDSC)-like states. In contrast, microglia were depleted in hypoxic areas, lacked compensatory CA isoenzymes, and developed TNF-linked stress responses and loss of homeostatic identity in acidic hypoxia. These findings identify metabolic adaptation to hypoxia-associated microenvironmental stress as a key determinant of GBM immune architecture, driving myeloid cell fates, spatial TME reorganization and the emergence of immunosuppressive tumor ecosystems.

---

IDH wild-type (IDHwt) glioblastomas (GBMs) are the most common malignant primary brain tumors in adults with a dismal median survival of only 15-18 months ^1^. GBMs are characterized by the profoundly immunosuppressive tumor microenvironment (TME), which is dominated by tumor-associated myeloid cells (TAMs), including brain-resident microglia (MG) and monocyte-derived macrophages (MDMs). TAMs not only support tumor progression but also actively suppress anti-tumor immunity and therapy responses ^2,3^.

Hypoxia is a defining feature of the TME in diffuse gliomas. It induces a series of adaptive mechanisms such as increased angiogenesis, tumor cell proliferation and invasion that all result in a more aggressive tumor phenotype and worsened patient prognosis ^4,5^. Hypoxia is also heavily linked to acidity, which is at least partially generated by the metabolic shifts in hypoxic conditions ^6^. In addition to directly influencing cancer cells, hypoxia has a profound effect on the tumor microenvironment ^7^. TAMs have been reported to display enhanced expression of immunoregulatory and proangiogenic genes in hypoxic niches, correlating with tumor aggressiveness and poor clinical outcome ^8–10^. Hypoxia also contributes to the compartmentalization of TAMs in GBM ^8,9,11^, yet the precise mechanisms underlying the spatial patterning of MDM and MG populations and their transcriptional states remain still poorly understood.

In this study, we applied a multi-modal strategy integrating cyclic immunohistochemistry (cIHC), single-cell RNA sequencing (SCS), spatial transcriptomics (ST), DNA methylation profiling, and *in vitro* cell cultures to characterize the impact of hypoxia on MDM and MG phenotypes in GBM. Our findings reveal a fundamental divergence in the hypoxia responses of MG and MDMs: MG undergo stress-induced dysfunction and depletion especially in the context of hypoxia-associated acidity, whereas MDMs adapt metabolically, survive, and acquire an immunosuppressive, myeloid-derived suppressor cell (MDSC)-like programs. These results uncover hypoxia and the associated acidity as central drivers that reorganize the myeloid landscape of GBM, offering mechanistic insight into macrophage dominance and MG exclusion within the tumor core. Together, this work provides a refined conceptual model of how microenvironment induced metabolic stress shapes tumor–immune interactions and the emergence of an immunosuppressive tumor ecosystem in GBM.

## Results

### Hypoxia mapping in diffuse astrocytomas identifies GBM-specific hypoxic niches

To define hypoxic regions and myeloid populations across diffuse astrocytomas, we performed cIHC for 173 IDH-wildtype GBMs and 49 IDH-mutant (IDHmut) astrocytomas (Fig. 1a, Table 1). Tumor samples were on tissue microarrays (TMAs) that included regions selected by a neuropathologist to ensure representation of vital tumor tissue. CA9, a canonical HIF1 target involved in intracellular pH homeostasis and extracellular acidification ^12,13^, served as an integrated marker of hypoxia and local acidosis. CA9 staining intensity, quantified by a QuPath-based pixel classifier, reliably distinguished normoxic, low-hypoxic, and high-hypoxic areas (Fig. 1a). To capture how hypoxia affects the myeloid compartment, we included only samples harboring at least low hypoxia for downstream analyses (136 GBMs and 25 IDHmut astrocytomas). High hypoxia occurred exclusively in GBM, whereas IDHmut samples contained only small, low-hypoxic foci based on CA9 staining (Supplementary Fig. 1a). In GBM, hypoxic domains were often extensive: over half of samples had ≥10% of cells residing in low (57%) or high hypoxia (52%). Hypoxic areas ranged from compact, viable pockets to larger regions bordered by pseudopalisading cells and micro-necrosis. Overall, the CA9 staining reliably highlighted the characteristic hypoxic regions in GBM, whereas in IDH-mutant tumors, it revealed only small, limited positive areas.

**Figure 1.**
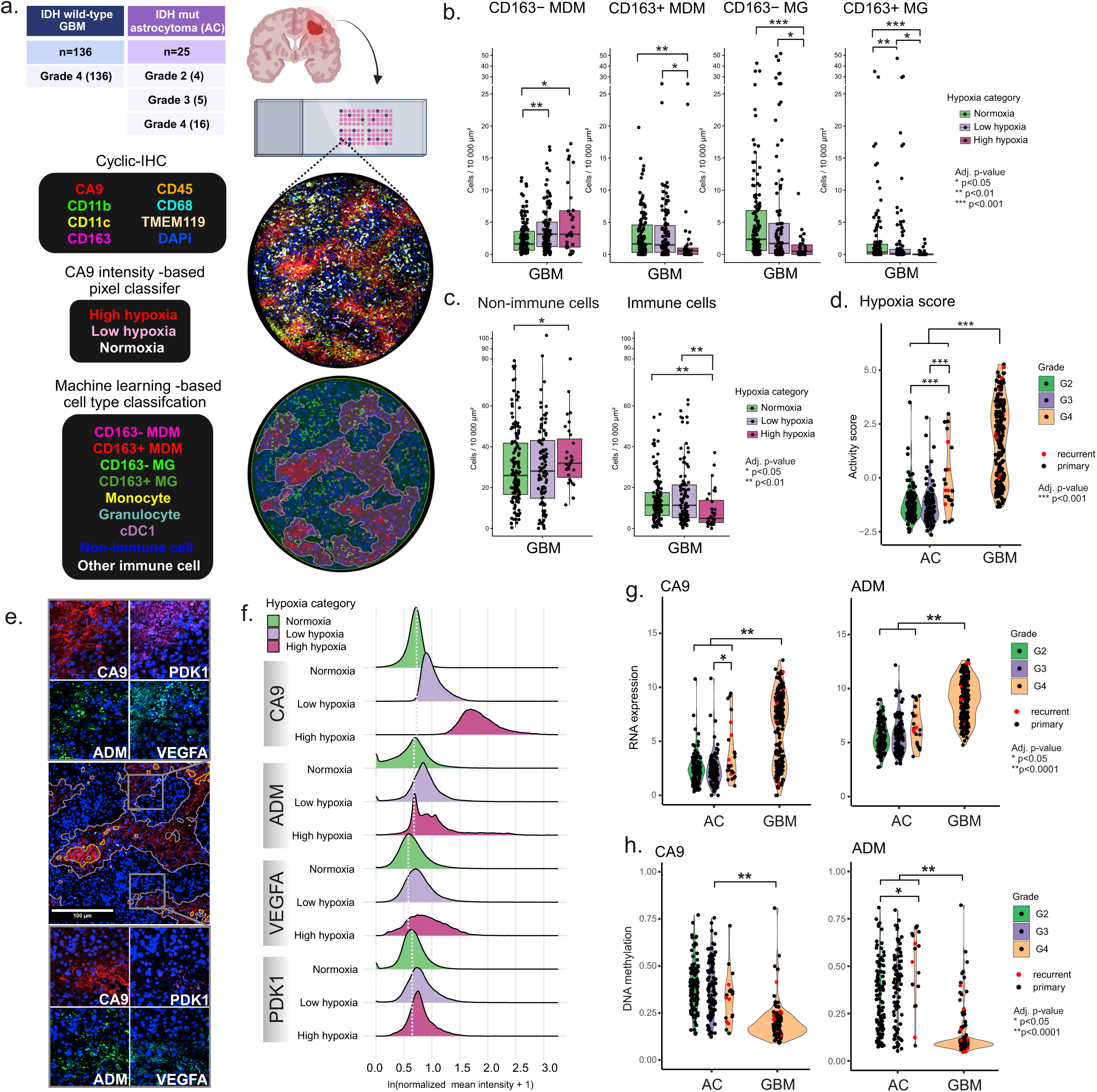
Hypoxia is associated with high-resolution spatial patterning of myeloid cells in GBM. a) Overview of the cIHC staining analysis, representing the used marker panel, image analysis, and sample cohort, created with Biorender ^14^. Cell types identified at single-cell level based on marker staining patterns and cell morphology were explored spatially in relation to CA9. b) MDM and MG cell type densities show significant hypoxia-associated differences in IDHwt GBMs. With an increase of hypoxia, the density of CD163-MDMs is increased, whereas CD163+ MDMs, CD163-MG, and CD163+ MG get depleted (Wilcoxon test). Normoxia n = 129, low hypoxia n = 101, high hypoxia n = 31. c) Densities of non-immune cells and immune cells across hypoxia categories in GBM cIHC data. Density of non-immune cells is significantly increased in high hypoxia whereas the overall density of immune cells is decreased (Wilcoxon test). Normoxia n = 129, low hypoxia n = 101, high hypoxia n = 31. d) Hypoxia score in 227 IDHmut astrocytomas (AC) and 203 IDHwt glioblastomas (GBM), calculated based on hypoxia response gene expression, is significantly higher in GBMs than IDHmut AC (diffuse astrocytoma) (Wilcoxon test). e) Representative image describing the typical staining patterns of all hypoxia markers (*CA9, ADM, PDK1 and VEGFA*) in a cIHC staining of GBM. ADM brightly positive cells co-accumulate with high CA9 positivity, VEGFA positivity concentrates on regions with low and high CA9 positivity, and PDK1 is only partially positive in the CA9 positive regions. f) Cell-wise quantified mean staining intensity of CA9, ADM, PDK1 and VEGFA in cIHC. Hypoxia categories were defined with a CA9-based pixel classifier. Ridge plots show relative staining intensity in respect to mean intensity of normoxia. In CA9-positive, hypoxic regions, the intensity distribution shifts towards right for all markers. Compared to CA9, the other three markers show a more narrow intensity range. g) Normalized expression of CA9 and ADM in 227 IDHmut AC and 203 IDHwt GBM are significantly higher in GBM than IDHmut ACs (Wilcoxon test). h) *CA9* and *ADM* show higher DNA methylation levels in IDHmut AC (n= 534) than GBM (n=155) (Wilcoxon test).

### Hypoxia drives selective accumulation of CD163□ macrophages and depletion of microglia in GBM

To look more in detail how different myeloid phenotypes are presented across different levels of hypoxia, we utilized machine-learning–assisted cell classification with cIHC staining data. The classification integrated six myeloid markers (CD11b, CD11c, CD45, CD68, CD163, TMEM119), nuclear morphology, and cell shape to identify four dominant myeloid subsets: CD163□ and CD163□ MDMs, and CD163□ and CD163□ MG along with rarer myeloid phenotypes, other immune cells, and non-immune cells (Fig. 1a; Supplementary Fig. 1b). Across GBMs, total immune cell density declined with increasing hypoxia (adj.p<0.01, Wilcoxon test), while non-immune cell density (representing predominantly malignant cells) increased (adj.p<0.05, Wilcoxon test) (Fig. 1c). Most distinct increase in cell density was identified for CD163□ MDMs, which increased in both low and high hypoxia (adj.p<0.001 and adj.p<0.05, Wilcoxon test, respectively; Fig. 1b). Conversely, both MG subsets and CD163□ MDMs were significantly depleted in high hypoxia (adj.p<0.01–0.001, Wilcoxon test; Fig. 1b). These patterns align with prior observations of macrophage core enrichment and microglial border predominance ^11^, but are here shown to be tightly linked to CA9-defined hypoxia. In grade 2-3 or grade 4 IDHmut astrocytomas, no significant hypoxia-dependent differences in MDM or MG abundance were detected (Supplementary Fig. 1c), likely reflecting the limited extent of CA9-associated hypoxia in these samples and possibly also a moderate sample cohort. Altogether, these findings demonstrate that GBM hypoxia specifically enriches CD163□ MDMs while excluding MG, revealing a selective ecological advantage for these cells in hypoxic niches.

### GBM exhibits stronger hypoxia response signatures and more permissive methylation constraints than IDHmut astrocytomas

To explore hypoxia response on transcriptomic level, we investigated the TCGA diffuse astrocytoma cohort (213 GBMs, 231 IDHmut astrocytomas). For this, we created a curated hypoxia-response gene set with correlation analysis that identified CA9, VEGFA, ADM, and PDK1 as the most robustly co-regulated hypoxia markers in these tumors (Pearson/Spearman >0.75; Supplementary Fig. 2a–b). A composite hypoxia score derived from these genes was significantly higher in GBMs than IDHmut astrocytomas (adj.p<0.001, Wilcoxon test; Fig. 1d). Among the four hypoxia response genes, CA9 exhibited the largest dynamic range with low activity in most IDHmut tumors but strong upregulation in GBMs (Fig. 1g) reflecting a similar pattern seen with cIHC. ADM, VEGFA, and PDK1 also showed similar but less pronounced trends in expression (Fig. 1g; Supplementary Fig. 2c). We confirmed a co-activation of these markers within CA9-positive regions with cIHC, which revealed spatial heterogeneity in their staining patterns (Fig. 1e-f). While bright ADM, VEGFA, and PDK1 were enriched in CA9-positive areas, the markers did not exhibit complete spatial co-localization. This suggests that hypoxia activates a shared program in GBM, but given the spatial heterogeneity, the degree of the response is likely cell type-related.

Since IDHmut tumors are known to exhibit global hypermethylation, we examined whether methylation constrains the expression of hypoxia response genes. CA9 and ADM promoters were significantly hypermethylated in IDHmut tumors compared with GBMs (adj.p<0.0001, Wilcoxon test) (Fig. 1h) and normal tissue (Supplementary Fig. 2f-g), correlating with lower gene expression (Supplementary Fig. 2d-e). VEGFA and PDK1 showed less differential methylation, consistent with their more comparable expression across tumor classes (Supplementary Fig.2c). These findings suggest that reduced hypoxia signaling in IDHmut astrocytomas is partly driven by gene hypermethylation, reinforcing the biological divergence between the two diffuse astrocytoma subclasses.

### Single-cell profiling delineates hypoxia-linked myeloid and malignant cell patterns in GBM

Next, we integrated four publicly available GBM single cell RNA-seq datasets (including 45 samples) to resolve hypoxia-associated myeloid states at single-cell resolution (Fig. 2a). Cell identities were assigned using canonical lineage markers, CD163 subsets based high/low CD163 expression and malignant cells were confirmed by inferCNV-derived chromosomal alterations (Fig. 2b). Hypoxia status was quantified using the *CA9–ADM–PDK1–VEGFA* gene set score, which displayed a trimodal distribution, allowing annotation of cells into normoxia, low hypoxia, and high hypoxia response classes (Fig. 2c). The distribution of myeloid subsets across hypoxia categories recapitulated our cIHC findings; CD163^low^ MDMs were strongly enriched in high hypoxia (57%), whereas MG of either CD163 phenotype had <5% of cells in this category (Fig. 2d). CD163^high^ MDMs exhibited an intermediate pattern, with 17% of cells in high hypoxia. Across malignant cell states, MES2-like, hypoxia-dependent GBM cells showed the highest degree of hypoxic response, as expected, although all the hypoxia categories were detectable across all tumor cell subtypes (Supplementary Fig. 3e). These patterns were independently validated using the GBMap dataset as reference atlas (Supplementary Fig. 3a–d). This allowed us to then connect the spatial patterns of hypoxia markers observed with cIHC to distinct cell-types. Expression of *ADM, VEGFA,* and *PDK1* increased progressively with hypoxia across all cell types, whereas CA9 induction was most pronounced in the malignant cells (Fig. 2e). The induction of hypoxia genes showed GBM cell subtype-associated patterns: strongest *VEGF* and ADM expression was detected in MES-like cells, CA9 in MES-like and NPC2-like cells, and PDK1 in NPC2 and OPC-like cells (Supplementary Fig. 3f). Among myeloid subsets, ADM was most strongly induced in CD163^low^ MDMs, consistent with their selective enrichment in cIHC-defined hypoxic regions (Fig. 2e, Fig. 1b). Taken together, the observed expression patterns of hypoxia response genes reflect the spatial heterogeneity seen in bright cIHC staining.

**Figure 2.**
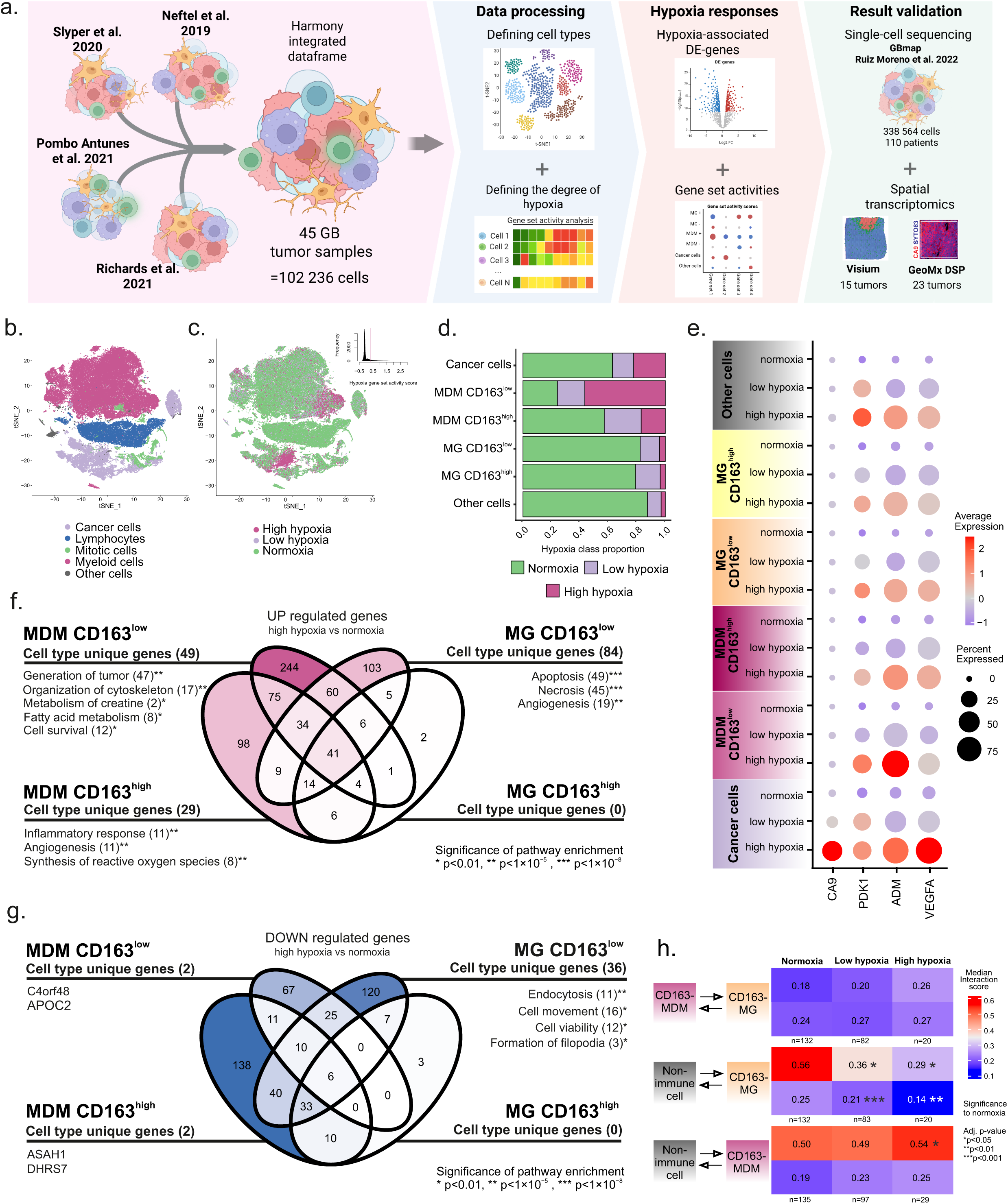
Myeloid subsets show cell-type dependent responses to hypoxia in single cell analysis. a) Concept figure of single-cell RNA-sequencing (SCS) data analysis, created with Biorender ^16^. The dataset was combined from four publications to cover 45 GBMs and utilized to study hypoxia responses at single-cell level. Cell types were identified using signature gene expression, and hypoxia using the response gene set. Differential expression (DE) analysis was done to identify cell type-dependent responses to hypoxia. Results were validated in external dataset ‘GBmap’ as well as GeoMx (37 ROIs from 23 tumors) and Visium ST (15 tumors, 46 012 spots) datasets. b) The t-SNE visualization of the SCS data set shows cell categories identified from the dataset. c) Distribution of hypoxia gene set activity scores in the t-SNE visualization of the SCS dataset. Activities were divided into normoxia, low hypoxia and high hypoxia based on thresholds seen in the histogram. The most prevalent hypoxic responses were detected in cancer cells and myeloid cells. d) Distribution of myeloid immune cell types into hypoxia categories. Majority of CD163^low^ MDMs are under high hypoxia, whereas most MG are under normoxia. e) Expression of hypoxia response genes across hypoxia categories in defined cell types in the SCS data. The highest CA9 expression is detected in cancer cells, whereas other hypoxia genes are upregulated in all the cell types. f-g) Venn diagram showing upregulated (f) and downregulated (g) gene counts in MG and MDM phenotypes in high hypoxia vs normoxia. Uniquely up- or downregulated genes by each cell type were run through IPA to identify processes they associate with. Uniquely up-/downregulated genes are both significantly up/downregulated by hypoxia and have significantly higher/lower expression in the specified cell type than in other cell types in high hypoxia, respectively. h) Interaction scores in cIHC data between non-immune cells, CD163-MDMs and CD163-MG. Cells with a centroid distance of <20 μm are considered interacting, and the interaction rate is normalized against the densities of respective cell types. In hypoxic regions, non-immune cells interact less with CD163-MG, whereas the interaction between non-immune cells and CD163-MDMs is increased.

### Hypoxia elicits divergent transcriptional and spatial responses in microglia and macrophages

Next, we sought to determine how hypoxia alters the transcriptional programs in MDMs and MG. We applied differential expression analysis (log2FC ≥ 0.25, adj. p ≤ 0.05) to SCS data, revealing both shared and lineage-specific responses. Across all myeloid subsets, only a small set of genes (42) was commonly upregulated, including primarily canonical hypoxia markers (*PDK1, ADM, VEGFA*), cytoskeletal stress markers (*CD44, VIM*), and the lipid-droplet–associated *PLIN2* (Fig. 2f).

CD163^low^ MG displayed the most extensive unique response, with 84 genes upregulated and 36 downregulated in high hypoxia. Upregulated genes were enriched for apoptosis, necrosis, and angiogenesis pathways (Fig. 2f), whereas the downregulated genes included key regulators of cell viability, cell movement, and endocytosis (Fig. 2g). Notably, 27 (55%) of the upregulated MG genes related to apoptosis were downstream targets of tumor necrosis factor (TNF), implicating it as a driver of the MG hypoxic response. Several downregulated genes belonged to the Arp2/3 complex, suggesting impaired cytoskeletal dynamics and reduced motility under hypoxia-induced stress.

In contrast, CD163^low^ MDMs upregulated 49 unique genes associated with cytoskeletal organization, migration, fatty-acid utilization, and cell survival (Fig. 2f), including *GATM* and *SLC6A8*, key components of creatine metabolism previously linked to macrophage fitness under hypoxia ^15^. Unlike MG, MDMs did not show upregulation of cell-death programs, indicating that MDMs remodel their metabolism to maintain viability, whereas MG undergo stress and metabolic failure in hypoxia.

Spatial interaction analysis further revealed lineage-specific adaptations to hypoxia. Under normoxia, non-immune cells were most frequently neighbored by CD163^−^ MG, followed by CD163^−^ MDMs. However, increasing hypoxia sharply reduced MG–non-immune cell proximity scores (adj.p<0.05, Wilcoxon test), while CD163^−^ MDM–non-immune cell interactions were significantly increased with hypoxia (adj.p<0.05, Wilcoxon test) (Fig. 2h). Residual CD163^−^ MG in hypoxia interacted more often with MDMs than non-immune cells, suggesting a reorganization of myeloid micro-architecture under metabolic stress.

### Hypoxia suppresses immune activity and promotes a macrophage trajectory toward immunosuppression

Many genes downregulated in CD163^high^ MDMs and MG were enriched for immune response and antigen presentation functions. To better capture these coordinated immune changes, we used gene set activity analysis to assess pathway-level responses. It revealed a progressive, hypoxia-associated decline in antigen presentation, interferon signaling, and leukocyte-activation pathways especially in CD163^high^ MDMs (Fig. 3a). Furthermore, CD163^low^ MDMs remained uniformly immune-inactive across all hypoxia states, but showed generally high activity and hypoxia-associated upregulation of previously reported early (E-MDSC) and monocytic (M-MDSC) myeloid-derived suppressor cell phenotype genes ^10^. To identify cell state transitions among TAM subsets and their associated molecular changes, we performed partition-based graph abstraction (PAGA) analysis (Fig. 3b). MDM population organized along a hypoxia-aligned pseudotime trajectory, which was not seen in MG (Supplementary Fig. 4b–c). Along this pseudotime trajectory (Fig. 3b–c), MDMs shifted from CD163^high^ to CD163^low^, which coincided with rising hypoxia score, increased activity of MDSC gene signatures along with cell survival genes characteristic of being upregulated by CD163^low^ MDMs in hypoxia. Furthermore, the activity of genes upregulated in both MDM phenotypes (n=75) was highest at the highly hypoxic end of the trajectory. In contrast, the immune-activity-related pathways (MHC-II, interferon response, phagocytosis) were progressively downregulated (Fig. 3c). These findings demonstrate that hypoxia not only selects for MDMs spatially but actively polarizes them toward immunosuppression.

**Figure 3.**
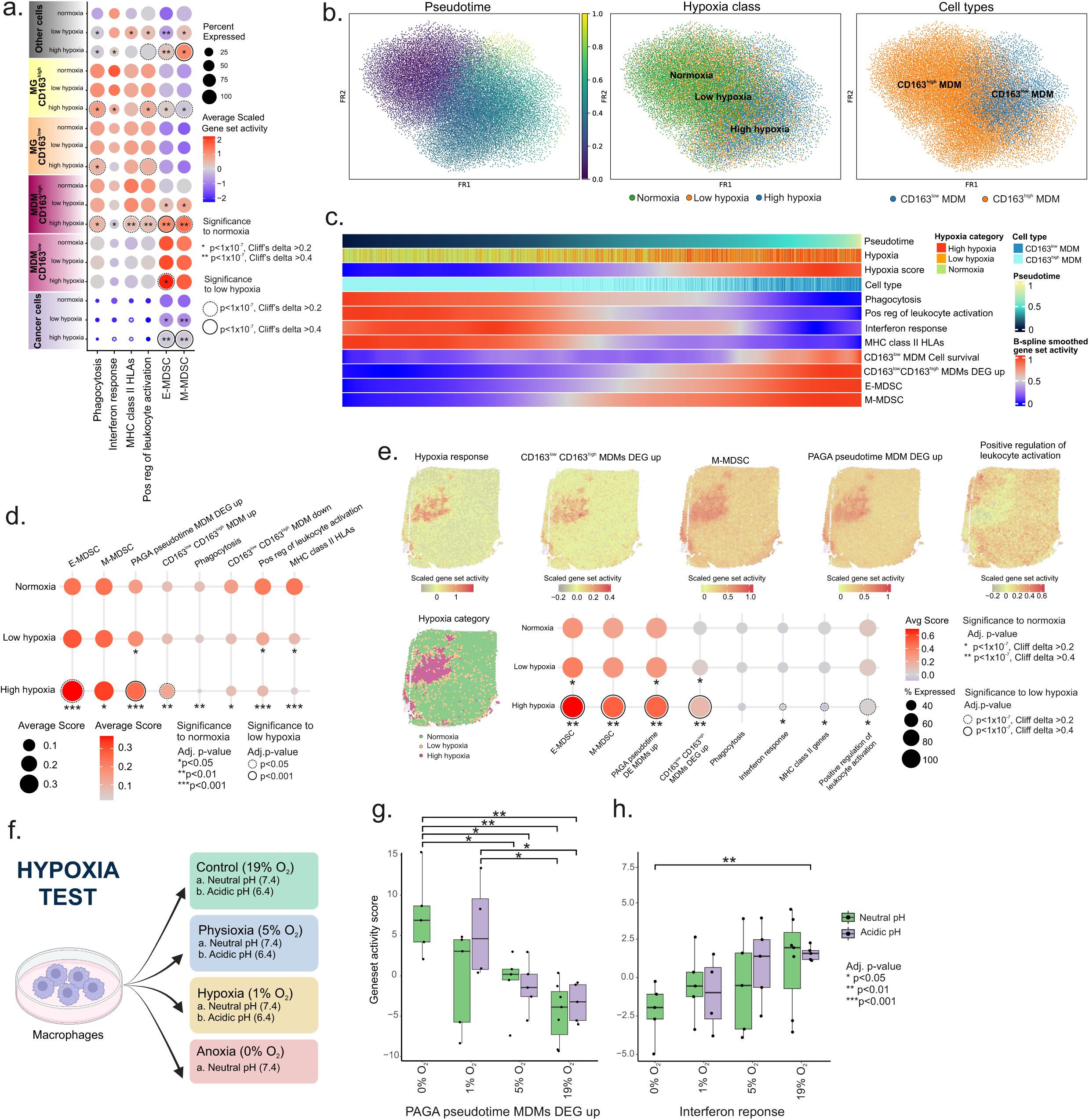
Hypoxia induces accumulation of immunosuppressive features in MDMs but not in microglia. a) Immune activity is downregulated in CD163^high^ MDMs, whereas CD163^low^ MDMs are overall immune-cold based on gene set activity analysis in SCS data. Interferon response is downregulated in MG and CD163^high^ MDMs. Wilcoxon test and Cliff’s delta was used for statistics to compare differences between different hypoxia groups within each cell type. b) PAGA trajectory analyses showing generated pseudotime trajectory and hypoxia categories in respect to MDM clusters. c) Gene set and reported MDM signature activities in PAGA trajectory support a shift of MDMs towards immunosuppressive phenotype with increasing hypoxia. CD163^low^ MDM Cell survival genes from IPA analysis, CD163^low^ CD163^high^ MDM DEG up genes and E- and M-MDSC signature genes from ^10^ are all upregulated along the pseudotime trajectory. Immune activity-related gene sets (Phagocytosis, positive regulation of leukocyte activation, interferon response and MHC class II) are all downregulated. d) In GeoMx spatial profiling data, immune activity-associated gene sets are downregulated in hypoxic regions, along with the increased activity of E-MDSC phenotype and MDM pseudotime trajectory genes upregulated in PAGA. e) Gene set activities in the 10x Visium dataset validate downregulation of immune activity and increase of E-MDSC, PAGA pseudotime MDM DEG up signatures with CD163^low^ CD163^high^ MDM DEG up genes (75 shared genes upregulated by both MDM subtypes) identified in SCS. f) Experimental setup to validate the direct effects of hypoxia and hypoxia-associated acidity on MDMs. PBMC-derived MDMs were cultured for 24 h in varying levels of oxygen tension with either neutral (pH 7.4) or acidic (pH 6.4) conditions. Image created with Biorender ^17^ .g) MDM PAGA trajectory genes are significantly upregulated in *in vitro* cell cultures when MDMs are exposed to extremely low oxygen tension (Wilcoxon test). h) Interferon response genes are significantly downregulated *in vitro* when MDMs are exposed to lowering oxygen tension (Wilcoxon test). For all conditions n=5, but in 1% O_2_ pH 6.4 n= 4.

To link single-cell level multi-modal observations to spatial context, we used both GeoMx DSP (37 ROIs from 23 GBM tumors, Supplementary Table 4) and 10x Visium datasets ^9^ (15 GBM tumors). Regions with high hypoxia-score (Fig. 3d-e; Supplementary Fig. 4e-f) showed marked downregulation of MHC class II, interferon response, and leukocyte-activation gene sets (Fig. 3d-e; Supplementary Fig. 4g). MDSC signatures and PAGA MDM pseudotime DE genes were strongly upregulated in hypoxic regions, providing spatial support for the pseudotime-derived, hypoxia-driven trajectory of macrophage suppression (Fig. 3d-e).

### Hypoxia alone induces MDM immunosuppressive features *in vitro*

To test whether hypoxia, alone or in combination with hypoxia-associated acidity, is sufficient to induce the identified transcriptomic changes, human peripheral blood mononuclear cell (PBMC)-derived macrophages were exposed to control (19% O□), physioxic (5% O□), hypoxic (1% O□), or anoxic (0% O□) environment with either neutral or acidic (pH 6.4) pH level (Fig. 3f). Transcriptomic profiling revealed significant upregulation of MDM pseudotime-trajectory genes especially in acidic 1% O□ and anoxic conditions (adj.p<0.05, adj.p<0.001, respectively, Wilcoxon test), whereas the interferon-response genes were gradually downregulated with decreasing O_2_ (adj.p<0.01, Wilcoxon test) (Fig. 3g). Lastly, acidity did not show a significant impact on gene set activities compared to the effect of decreased oxygen levels alone (Fig.3g-h, S5a). Altogether, our results suggest that severe oxygen depletion directly promotes the transition of macrophages to the suppressive phenotype trajectory observed *in vivo*.

### MG mount a TNF–driven stress response under hypoxia, leading to vulnerability and dysfunction

As MG responses to hypoxia were associated with compromised survival (Fig. 2f-g) and TNF-induced responses (Supplementary Table 3), we ran gene set activity analysis to further characterize cellular responses in MG. Our results highlighted TNF and NF-κB mediated activation being the most prominent in CD163^low^ MG under hypoxia (Fig. 4a). This was accompanied by high activity of stress-response genes in these cells. Furthermore, TNF was more strongly expressed in MG than MDM, and TNF receptor 1 (*TNFR1*, alias *TNFRSF1A*) and 2 (*TNFR2*, alias *TNFRSF2A*) were also clearly expressed in MG (Fig. 4a). TNFRSF1A acts as the main mediator of TNF-induced cell death and an activator of NF-κB signalling^18^. CD163^low^ MDMs showed low expression of TNF and both receptors especially in highly hypoxic conditions. Stress response, TNF- and TNF–NF-kB responses were all also significantly upregulated spatially in hypoxic regions (10x Visium) (Fig. 4b). *In vitro*, microglia exposed to acidity and hypoxia reproduced these patterns (Fig. 4c-e, Supplementary Fig. 5a-b). TNF–NF-kB signaling was significantly upregulated already in acidic physioxia (adj.p<0.01, Wilcoxon test) (Fig. 4d). Consistently, TNF was significantly upregulated in MG in physioxia and hypoxia under acidic conditions (adj.p<0.05, Wilcoxon test) (Fig. 4e). Additionally, TNF receptors had an overall higher expression in MG than MDMs (Fig. 4e).

**Figure 4.**
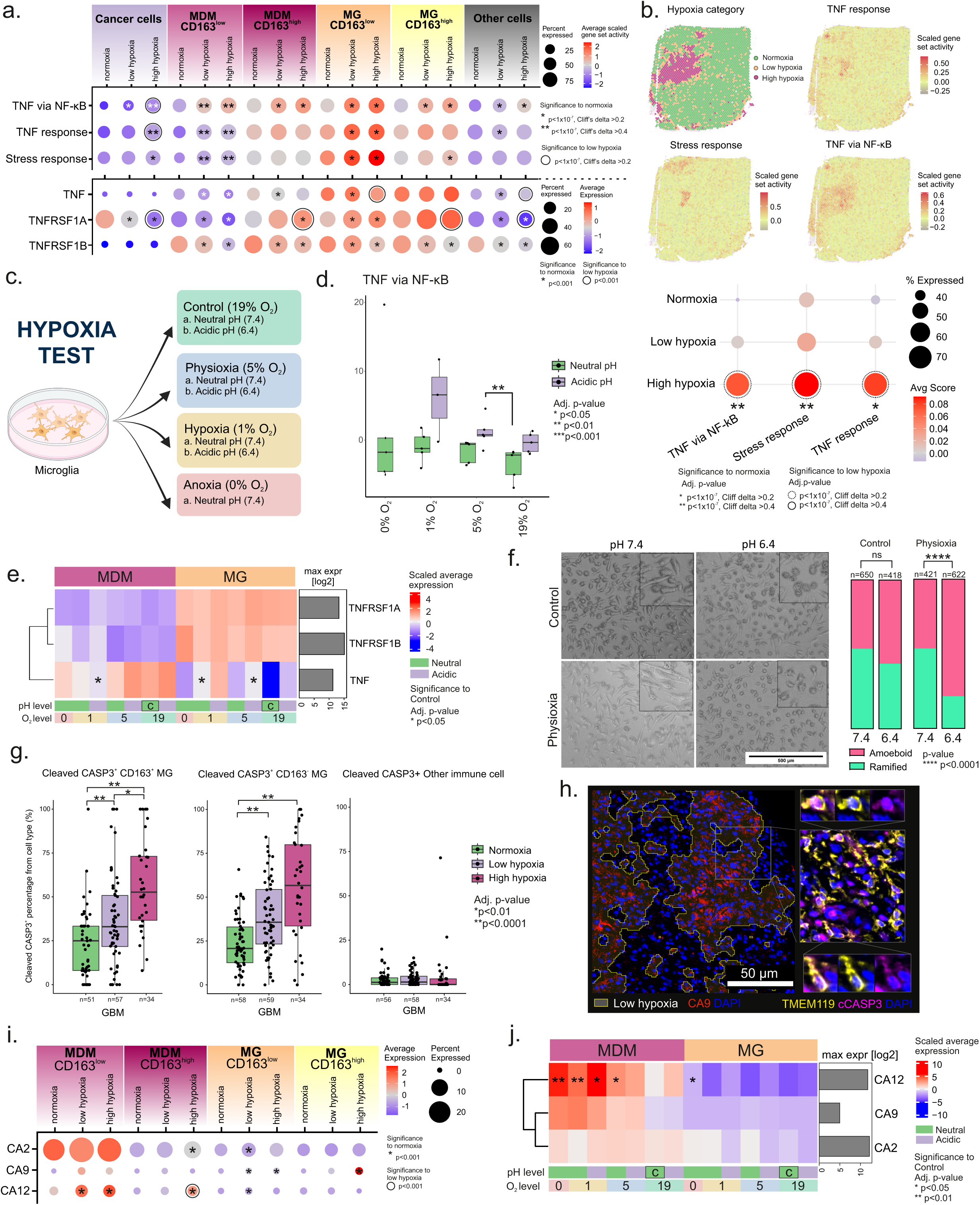
Microglia are characterized by TNF-signalling related stress signatures and heightened sensitivity to hypoxia-associated acidity. a) Highest activities of stress, TNF, and NF-kB-mediated TNF response gene sets are detected in CD163^low^ MG along with elevation in TNF and its receptors 1 (TNFRSF1A) and 2 (TNFRSF1B). Wilcoxon test, with Cliff’s delta for gene sets, used for statistics to compare differences between hypoxia groups within each cell type. b) TNF, NF-kB-mediated TNF, and stress response genes show significant upregulation in hypoxic regions in the 10x Visium dataset. c) Hypoxia test setup to validate MG responses *in vitro*. Human induced pluripotent stem cell (hiPSC)-derived MG were cultured for 24 h in varying levels of oxygen tension with either neutral (pH 7.4) or acidic (pH 6.4) conditions. Image created with Biorender ^23^ .d) Activity of TNF signaling via NF-kB in *in vitro*-cultured MG shows significant upregulation in acidic physioxia and a general elevation in acidic culture conditions (Wilcoxon test). For all conditions n=5, but in 0% O2 pH 7.4 n= 4 and 1% O2 pH 6.4 n= 3. e) Scaled (log2) expressions of TNF and its receptors TNFRSF1A and TNFRSF1B in *in vitro*-cultured MDMs and MG. TNF is significantly upregulated in acidic hypoxia in MG, whereas decreased expression is detected in MDMs compared to control conditions. Expression of the TNF receptors is higher in MG compared to MDMs (Wilcoxon test). f) Microglia adopt an amoeboid shape under acidic pH in physioxia after 48h (Fisher’s exact test). g) Apoptotic signaling activity, namely cleaved caspase 3 (cCASP3) positivity, is dramatically increased in MG with increasing degree of hypoxia (Wilcoxon test), aligning with their compromised viability. h) Representative cIHC staining of cCASP3+ MG (detected with TMEM119) (on the right) in a low hypoxia area in a GBM sample (on the left). i) Strongest CA2 expression and hypoxia-induced CA12 upregulation (Wilcoxon test) is detected in CD163^low^ MDMs in the SCS data. j) *In vitro*-cultures of MDMs and MG validate the differential expression patterns of CA2, CA9 and CA12 with highest expression in anoxic and hypoxic MDMs (Wilcoxon test).

To further observe the effect of acidity on MG, cells were exposed to lowering pH in either physioxia or control oxygen level for 48 hours (Supplemental Fig. 5c). The cells displayed marked morphological changes from ramified to rounded, amoeboid forms (Fig. 4f; Supplementary Fig. 5d-e). Acidic conditions combined with decreased oxygen produced the most pronounced morphological distress, which is in line with identified downregulated processes for MG in SCS data, such as the downregulation of formation of filopodia (Fig. 2g). Furthermore, MG cell numbers also modestly decreased with the decrease of pH level and oxygen tension (Supplementary Fig. 5f), consistent with the transcriptional stress signatures. Oxygen-gradient assays (Supplementary Fig. 5g) showed no directed migration away from hypoxic zones (Supplementary Fig. 5h), indicating intrinsic susceptibility rather than escape behavior. The impaired viability of MG in CA9-associated hypoxic regions in GBM was verified with cIHC, which showed increased fraction of Cleaved caspase −3 (cCASP-3) positive MG in low and high hypoxia regions (adj.p<0.0001, Wilcoxon test) (Figure 4g-h). Furthermore, the intensity of cCASP-3 increased in both MG subtypes with the increase of hypoxia (Supplementary Fig. 5i).

Altogether, anoxia did not induce as intense TNF, NF-κB, and stress responses as low oxygen levels combined with added acidity in cultured MG. In addition to transcriptional level, the cells also experienced morphological stress in acidic conditions *in vitro*. Our results suggest that MG, unlike MDMs, experience stress especially in hypoxia-associated acidity.

### Divergent carbonic anhydrase expression explains the differential resilience of MDM versus MG

To further explain the different capabilities of MDMs and MG to survive environmental stress, we looked into their expression of carbonic anhydrases, the enzymes that are the master regulators of acid-base homeostasis ^19,20^. Single-cell analysis showed that CD163^low^ MDMs express higher levels of CA2 and CA12 compared to MG, and even CA9 positivity was detected in CD163^low^ MDMs under hypoxia (Fig. 4i). These results were validated *in vitro*, along with hypoxia-induced CA12 upregulation (Fig. 4j). CA12, CA9, and CA2 form a pH-regulating system in which CA2 functions in the cytosol, while membrane-bound CA9 and CA12 catalyze extracellular acidification ^20^. Together, these enzymes support metabolic adaptation and survival ^21,22^. The observed differential CA expression likely governs intracellular pH buffering capacity, conferring macrophage resilience and microglial vulnerability in acidic hypoxia.

## Discussion

We integrated cyclic immunohistochemistry, single-cell RNA sequencing, spatial transcriptomics datasets, DNA methylation profiling and *in vitro* cell cultures to characterize the impact of hypoxia and hypoxia-associated acidity on MDM and MG phenotypes in GBM. Our analysis reveals that CA9-associated hypoxia and the resulting acidity in the tumor microenvironment have distinct effects on MDMs and MG. Based on our results, MDMs acquire immunosuppressive phenotype and MDSC-like programs under hypoxic stress, while their fitness in the harsh environment is being promoted by CA2 and CA12 expression. Consistent with our findings, CA12 expression has been linked to macrophage survival in an acidic TME in other cancer contexts ^24^, suggesting a conserved mechanism of myeloid adaptation to low-pH niches. The survival of the MDMs also appeared to be supported by metabolic reprogramming involving fatty acid utilization and creatine, in line with previous findings ^15,25^.

In contrast to MDMs, MG exhibit heightened sensitivity to hypoxia-associated pH changes. Combined hypoxia and acidity induce TNF-linked cellular stress and reduced viability. Unlike MDMs, MG show little evidence of metabolic adaptation or carbonic anhydrase upregulation, suggesting a restricted capacity to withstand the acidic hypoxic conditions. At the tissue level, these cell-intrinsic differences translate into reorganized spatial interaction patterns. Under hypoxia, MG-tumor cell engagement is dramatically decreased, indicating a reorganization of cellular networks within the TME. These phenotypic and interaction changes drastically modulate the cellular landscape and neighborhoods in the GBM tissue upon oxygen depletion and acidification.

Analyses in IDH-mutant tumors were limited by their low prevalence in our cohort. Within these constraints, we did not detect similar hypoxia-associated changes in myeloid cell densities comparable to those seen in GBM. Although necrosis, heavily linked to severe hypoxia, is present in part of the grade 4 IDHmut tumors, necrotic regions tend to be more focal and present in a smaller fraction of tumors ^26^. This was also supported by our analysis, detecting mainly small CA9-associated hypoxic regions in IDHmut astrocytoma samples, aligning with previously reported characteristics ^27^. Vascular abnormalities also appear less prevalent in grade 4 IDHmut astrocytoma than in GBM ^26,28^, coinciding with lower ADM activity in our datasets. Mechanistically, our analysis suggests that IDH mutation and associated DNA hypermethylation suppress *CA9* and *ADM* activity, implying downstream consequences also for the microenvironment. A similar epigenetic repression mechanism has been described for HIF-1 target genes involved with pH regulation, *e.g.* lactate dehydrogenase A (*LDHA*), in IDHmut diffuse astrocytomas ^29^, supporting the broader view that IDH-driven epigenetic programs blunt hypoxia adaptations.

Collectively, our findings highlight the importance of distinguishing between brain-resident microglia and monocyte-derived macrophages, as their distinct responses to hypoxia and acidity reveal cell-type-specific opportunities for therapeutic intervention. Moreover, both oxygen tension and pH should be taken into account when assessing cellular behaviour within the tumor microenvironment, as these factors impose complementary and at times divergent pressures on immune populations. Our results further highlight the contribution of CA isoenzymes to myeloid cell persistence within malignant brain tumors.

Recent work has shown that hypoxia□driven TAM subsets can destabilize endothelial junctions through ADM secretion, and that therapeutic blockade of this axis can restore vascular integrity and enhance drug delivery in GBM models ^30^. Together with our observations, these findings suggest that targeting hypoxia□induced signaling pathways, such as carbonic anhydrase–mediated pH buffering ^31^, may offer complementary avenues for TME reprogramming. Such strategies have the potential to alleviate immunosuppression, and ultimately improve therapeutic efficacy in GBM.

### Supplementary material

**Supplementary Table 1.** Tumor characteristics of cIHC study cohort.

**Supplementary Table 2.** Gene sets and differentially expressed genes. List of genes included in the analyzed gene sets and those that are differentially expressed in high hypoxia when compared to normoxia in the SCS analysis.

**Supplementary Table 3.** Ingenuity pathway analysis (IPA) results of DE genes for MDM and MG subclasses.

**Supplementary Table 4.** GeoMxDSP counts and metadata.

## Online methods

### cIHC analysis cohort

The cohort consists of 222 diffuse astrocytoma samples. Samples with at least low hypoxia, and a minimum of 20 cells in that region were then included for the analyses resulting in 161 samples in total. Samples previously identified as low-grade IDHwt (before the year 2020) were re-classified as grade 4 GBs (n=5), as long as the patient had been over 20 years old. Low-grade IDHwt samples from patients under 20 years old were omitted from the analysis. The sample material was formalin-fixed paraffin-embedded (FFPE) tumors (fixed with 4% phosphate-buffered formaldehyde) sampled into 7 tissue microarrays. An experienced neuropathologist evaluated the original whole-mount FFPE tumor samples and determined the histopathological type and grade according to the criteria presented by the World Health Organization (WHO).

### cIHC

The 4 μm thick sections were stained with in-house cIHC protocol based on Multiple Iterative Labeling by Antibody Neodeposition (MILAN)^35^. Antigen retrieval was done using a Tris-HCl buffer (pH 9.0) prior to antibody labeling in 121□ for 20 min. Non-specific epitopes were blocked using normal goat serum (S-1000, Vector Laboratories) 1:20 for 5 min in room temperature and autofluorescence was treated with 0,1% Sudan Black solution (199664, Sigma-Aldrich) for 5 min in room temperature. Myeloid immune cells and hypoxia were stained with anti-CD11b (D6X1N, Cell signaling technology, #49420S) 1:100, anti-CD11c (D3V1E, Cell signaling technology, #45581S) 1:200, anti-CD163 (10D6, Leica Biosystems, CD163-L-CE) 1:100, anti-CD68 (EPR20545, Abcam, ab213363) 1:400, anti-Ca9 (M75, Atlas antibodies) 1:50, anti-CD45 (D9M8I, Cell signaling technology, #13917S) 1:200, anti-TMEM119 (polyclonal, Prestige Antibodies, #HPA051870) 1:1000. Hypoxia response marker panel was stained using anti-PDK1 (Proteintech, #18262-1-AP) 1:300, anti-VEGF (JH121, Invitrogen, MA5-13182) 1:50, anti-Ca9 (M75, Absolute antibodies, #Ab00414) 1:50 and anti-ADM (R&D Systems, 1003019, #MAB61081-SP) 1:50. Apoptotic cells were stained using anti-Cleaved caspase-3 (Asp175, Cell Signaling Technology, #9661)1:200, with aforementioned CA9, TMEM119, CD45, and CD163. Detection was done with fluorescently labeled secondary antibodies (Donkey anti-Mouse IgG (H+L) Highly Cross-Adsorbed Secondary Antibody, Alexa Fluor 647 (Invitrogen, A31571), Goat anti-Rabbit IgG (H+L) Highly Cross-Adsorbed Secondary Antibody, Alexa Fluor Plus 647 (Invitrogen, A32733), Goat anti-Mouse IgG (H+L) Cross-Adsorbed Secondary Antibody, Alexa Fluor 750 (Invitrogen, A21037)). Samples were mounted with Fluoromount-G (Invitrogen, 00-4959-52) mounting medium that contained DAPI for tissue counterstain. The staining result was scanned using NanoZoomer S60 (Hamamatsu) whole-slide scanner using 387 nm, 650 nm and 740 nm excitation wavelengths. The fluorescent cIHC staining was followed by an HE-staining on the same tissue section. HE-staining was done using KEDEE KD-RS3 Automatic Slide Stainer and the slides were mounted using DAKO coverslipper with DPX mountant (Sigma-Aldrich, 44581).

### Image registration

Registration of whole slide images was achieved by taking all DAPI images from different staining rounds and assigning one of them as the fixed reference. All the other DAPI images were then aligned to it in a pairwise manner. Registration of an image pair started by finding SIFT features from an image that has its long edge subsampled to 2048 pixels while maintaining aspect ratio. Matching features between images were found by Fast Library for Approximate Nearest Neighbors (FLANN) and transformation parameters are estimated using Random Sample Consensus (RANSAC). The found affine transform was then fine-tuned by minimizing Normalized Gradient Fields (NGF) distance between the images, which aims to match orientations of gradients for the two images making it suitable for multi-stain registration ^36^. This registration process was repeated for three different values of epsilon (10, 60, 110). The best of all three registrations was selected by visual inspection. In case automatic registration did not produce sufficient accuracy, images were annotated manually for 3 landmark points and an optimal affine transform in the least squares sense was applied. Insufficient registration accuracy was also detected via visual inspection. The transformations acquired for each DAPI image, whether obtained by automatic or manual approach, were then applied also to the marker images to obtain registered TMA stacks.

Fiji ^37^ was used for fixing the registering of individual images, utilizing the TrakEM2 plugin ^38^. Images were first loaded into a new TrakEM2 project by importing the image stack. The images selected for registration were aligned using the *Align Layers* feature on the selected layers and choosing the appropriate alignment affine method. Alignment parameters, including the number of iterations and convergence threshold, were adjusted as needed.

In addition, the 2nd cIHC staining set and hypoxia marker cIHC staining set images were registered with VALIS ^39^ (v1.2.0), utilizing the DAPI of the first round of staining as the fixed reference. The DAPI stainings underwent registration, and after that, the changes were applied to the marker images via transformation.

### cIHC Image analysis

The images were analyzed using ImageJ 2.3.0/1.53q / Java 1.8.0_172 and QuPath v0.4.3. The registered images were processed into hyperstacks using a custom-written ImageJ Macro and the hyperstacks were loaded into QuPath. Cells were detected using the automated cell detection function based on the DAPI channel. To avoid including necrotic cells in the analysis, the cell min area, median filter, sigma and threshold values were adjusted from default parameters. Larger necrotic regions were excluded from the analyzed area. Hypoxia was classified into three categories based on CA9 staining intensity. Areas were annotated with a pixel classifier using intensity threshold for high hypoxia (>10), low hypoxia (4≤ x ≤10) and normoxia (<4).

Cells were classified with a machine learning-based (Random Trees) object classifier. Classifier was trained with input of intensity features of all markers (1st cIHC set: CD11b, CD11c, CD45, CD68, CD163, TMEM119, DAPI / 2nd cIHC set: CD45, CA9, TMEM119, cCASP3, CD163, DAPI) measured from cell, nucleus and cytoplasm area (min, max, mean and st. dev), smoothed features with radius (FWHM) of 25 and 75 pixels and Haralick features. The classifier was trained by manually annotating cells to predetermined classes. Microglia annotation was based on TMEM119+, and further divided to CD163- and CD163+. In the first set microglia were also CD11b+, CD11c+, CD45+, and CD68+ with varying intensities, but this information was not used as criteria for identifying the cells. Macrophages were identified with CD68+, CD45+, and CD11b+ and TMEM119-. Macrophages showed varying intensity of CD11c, but it was not used as a phenotyping criterion. cDC1 class was annotated with CD11b+ and CD11c+, and CD68-, CD163- and TMEM119-. Monocytes and granulocytes were positive only for CD11b and CD45 and the two cell types were separated based on cell size and shape of the nuclei. ‘Other immune cell’ class was positive only for CD45. Cells negative for all markers were annotated to the ‘Other cell’ class. CASP3 was used as an additional label in the 2nd cIHC analysis set to divide cells into groups based on its positivity. Each phenotype was given a minimum of 50 example cells to train the classifier. Intensities of hypoxia markers (CA9, ADM, VEGFA, PDK1) were measured from the whole estimated cell area based on cell segmentation that was done using DAPI staining.

Sample-wise marker intensity values (CA9, VEGFA, ADM, PDK1, cCASP3) were normalized to the mean intensity value in the nonhypoxic area.

### Statistical analysis of the cIHC data

Statistical analyses for cIHC data were performed using R (v4.4.1). The densities and fractions of all cell types were calculated in respect to the sample’s hypoxia regions. Small areas of normoxia, or low or high hypoxia (< 20 cells in total for the sample) were omitted from the analysis. The extent of hypoxic regions was quantified by calculating the percentage of pixels in each TMA core covered by hypoxia.

### TCGA data analysis

TCGA gene expression and methylation data were obtained from the GDC legacy database and reclassified based on WHO 2021 classification as previously described ^40^. Methylation probes annotated to the genes of interest were filtered based on two criteria: maximum distance to the gene transcription start site of <10,000 bp and variance >0.02 across all TCGA methylation samples (155 glioblastoma and 534 low-grade glioma cases). For CA9 and ADM genes, the maximum distance between filtered probes and transcription start sites were 2440 and 2243 bp, respectively. Median beta value of filtered probes was used to represent gene methylation after inspecting correlation with gene expression. Significant differences in log2-transformed RNA expression (595 TCGA cases) or DNA methylation (496 TCGA cases) between glioma subtypes or tumor grades were calculated using the Wilcoxon rank-sum test. Hypoxia gene set was created and activity z-score calculated in TCGA diffuse astrocytomas similarly as in ^41^. Heatmaps, violin plots, and scatter plots from TCGA data were created using R (v4.0.4) and packages ggplot2 (v3.3.5), gplots (v 3.1.3.1), and ComplexHeatmap (v 2.6.2).

### Methylation analysis with a wider cohort

The raw data from ^33^ was downloaded from GEO (GSE90496) and preprocessed and normalized similarly as in ^33^, including the material batch effect correction (FFPE/Frozen) and following probe filters: 1) removal of probes targeting the sex chromosomes, 2) removal of probes containing a single-nucleotide polymorphism (dbSNP132 Common) within five base pairs of and including the targeted CpG site, 3) probes not mapping uniquely to the human reference genome (hg19) allowing for one mismatch, and 4) probes not included on the Illumina EPIC array ^42^. Median probe values were visualized with ggplot2 (v3.3.5). The same promoter region DNA methylation probes annotated to the genes of interest (*CA9*, *ADM*) were used as in the TCGA analysis.

### Cell interaction analysis

Cell interaction analysis was performed for the cIHC data using *cKDTree* (Scipy v1.11.3) in Python (v3.9.18), using the method described in ^43^. The distance between cells was set to be a maximum of 20 μm, after tuning and testing out different distances. To keep the data consistent with the statistical analysis, the small areas of hypoxia (<20 cells in total for the sample) were omitted. Results were analysed in R (v4.4.1).

### Single-cell RNA sequencing data analysis

Single-cell RNA sequencing count matrices from GB samples were obtained from ^3,44–46^ and preprocessed in R (v4.3.0) to contain cells with expression from a minimum of 200 genes and genes with expression in a minimum of 3 cells. The threshold for mitochondrial gene expression was set to 5% for all cells. After filtering, 102 236 cells from 45 GBMs were retained for analysis. Copy number alterations were referred using gene expression means with a window length of 301 by inferCNV (v1.10.1)^47^. Cells were organized into 5 (data from ^45,46^) or 10 (data from ^44^) clusters and copy number alterations of chromosomes 7 and 10 were used to identify copy number altered clusters. Data from ^3^ was CD45+ prefiltered, and thus did not undergo copy number alteration analysis.

Data was integrated into a Seurat object (v4.3.0.1) ^48^ using Harmony (v1.2.0) ^49^ with 2000 variable features, 50 PCs and original identities as the grouping variable. The cells were organized into clusters using Seurat’s *FindNeighbors* (40 dimensions) and *FindClusters* functions (resolution 3). Cell types were annotated manually for each cluster using the expression of known cell type marker genes. Cancer cells were identified using the copy number variation analysis results and their subtypes based on gene signature ^3,44–46^ activity scores calculated with Seurat’s *AddModuleScore* function similarly as in ^44^. The data was visualized with the tSNE dimensional reduction technique using Seurat’s function *RunTSNE*.

The differentially expressed genes were obtained using Seurat’s *findMarkers* function with MAST (v1.28.0). A logarithmic fold change threshold was set to 0.25 and a Bonferroni-corrected p-value to 0.05. Pathway analyses were done using Ingenuity Pathway Analysis software (Qiagen).

### Single-cell gene set activity analysis

Gene sets were collected from The Molecular Signatures Database (MSigDB) and selected publications ^10,50–52^. Gene sets were filtered based on positive pair-wise correlation between the genes in TCGA IDHmut astrocytoma and GB (see below) expression data. The correlations were first clustered and within the resulting clusters of 4-20 genes, seed genes correlating with each other with Pearson and Spearman correlation coefficients >0.4 were drawn and other genes correlating with the seed genes (>0.4 Pearson and Spearman) were added to the final filtered gene set.

All gene set scores were calculated for each cell using Seurat’s *AddModuleScore* function. For scoring the cells into hypoxia categories, the hypoxia geneset (*CA9, VEGFA, PDK1, ADM*) was utilized, and based on the scores, the data was divided into three groups: normoxia (<0), low hypoxia (0-0.3), and high hypoxia (>0.3).

### PAGA Trajectory analysis

For the microglia and macrophage populations in the single-cell RNA-sequencing data, pseudotimes were inferred using partition-based graph abstraction (PAGA) implemented in Scanpy (v1.9.6) with Python (v3.9.18). During initial clustering and PAGA analysis of the microglia subset, a small outlier cluster originating mostly from one sample (Cluster 13, Supplementary Fig. 3a) was identified and removed before re-running PAGA. The pseudotime starting point for each cell type was selected as the cell with the lowest hypoxia gene set activity score: a CD163^high^ macrophage for the macrophage subset and a CD163^low^ microglia for the microglia subset.

Differentially expressed genes along pseudotime in the macrophage population were identified by fitting a linear model of gene expression as a function of pseudotime (R v4.4.1). Gene set activity scores were smoothed using the *smooth.spline* function from the base R stats package with default settings.

### GeoMx Digital Spatial Profiling

The analyzed samples were selected through quality control experiments done with RNAScope Multiplex Fluorescent Reagent Kit v2 (ACDBio, Cat #323135). Assay was imaged with whole-slide fluorescent imaging (Olympus VS200). Samples with positivity for all three control genes (PPIB, POLR2A, UCB) were accepted for the GeoMx DSP cohort. Morphology staining was first optimized with hypoxic GB tissue. Hypoxia was stained using recombinant Alexa Fluor 647 Anti-Carbonic Anhydrase 9/CA9 antibody 1:50 (EPR4151(2), ab225074) and nuclei with SYTO83 Orange Fluorescent Nucleic Acid Stain (S11364, Invitrogen). Regions of interest (ROIs) were selected based on the morphology staining. Areas with high CA9 positivity, low CA9 positivity and CA9 negative areas were selected and annotated based on visual evaluation of staining intensity into normoxia, low hypoxia or high hypoxia categories. For data-analysis, 4 out of 54 total ROIs were then annotated into another class based on hypoxia gene set activity for more precise categorization due to misinterpretation of CA9 staining. In the ROIs, autofluorescence and especially high vascularization caused the misclassification. The sequencing library was prepared from collected areas of interest according to the manufacturer’s protocol (MAN-10117-05).

### GeoMx data analysis

Fastq files were processed through the GeoMx NGS Pipeline Linux (v3.1.1.6) using default settings. The results were analyzed in R (v4.4.1) using the NanoStringNCTools (v1.12.0), GeomxTools (v3.8.0) and GeoMxWorkflows (v1.10.0) libraries. For each ROI to be included, the QC parameters were set to *minSegmentReads = 1000, percentTrimmed = 80, percentStitched = 80, percentAligned = 80, percentSaturation = 60, minNegativeCount = 1, maxNTCCount = 9000, minNuclei = −1, minArea = 2500*. qcCutoffs were set to *minProbeRatio = 0.1, percentFailGrubbs = 20* with *removeLocalOutliers = TRUE*. Limit of Quantification (LOQ) cutoff and minimum LOQ were both set to 2. Gene detection threshold and segment cutoff were set to 5%. Data was then Q3-normalized. Gene set activities were calculated using AUCell (v1.28.0). Hypoxia categories were assigned to the ROIs based on the hypoxia geneset score. ROIs with gene set score < 0.15 were assigned as normoxic, 0.15 - 0.30 as lowly hypoxic, and > 0.30 as highly hypoxic.

### Visium

GMB tumor core samples from ^9^ were re-analysed using Seurat (v5.3.0). The data was filtered to contain only spots with > 500 genes, and genes were required to be expressed in at least 5 spots. One sample (UKF265_T_ST) was excluded from the analysis due to insufficient spots remaining after filtering, resulting in 15 samples and 46 012 spots for downstream analysis. The samples were individually normalized using the Seurat *SCTransform*, after which gene set analysis was performed using Seurat *AddModuleScore*. Spots were assigned to hypoxia classes based on the hypoxia geneset score, with those < 0 being normoxic, 0 - 0.24 lowly hypoxic and > 0.24 highly hypoxic. In addition to p-value testing, Cliff’s Delta values were calculated as in the SCS analysis to estimate the effect size change.

### PBMC-derived macrophages

Whole-blood samples were collected from several healthy individuals. Both sexes were covered. Buffy coats were isolated using Ficoll Paque PLUS (GE17-1440-03, Cytiva). Monocytes were then isolated through adherence-selection using Monocyte Attachment Medium (Promocell, cat. C-28051) and differentiated using M2-Macrophage Generation medium XF (Promocell, cat.C-28056) according to the manufacturer’s protocol.

### Human iPSC-derived microglia

HiPSC line UTA.04511.WTs was used for the differentiation of microglia-like cells according to previously published protocol with minor modifications ^53,54^. On day 16, cells were seeded on tissue-culture treated 96-well plates at 15,000 cells/well or to hypoxia chambers (BioGenium Microsystems Oy) that were coated using poly-L-ornithine (100 µg/ml) (Sigma Aldrich, P3655) and recombinant human laminin-521 (30 µg/ml) (BioLamina, LN521) at a density of 50,000 cells per chamber. Cells were exposed to hypoxia and/or acidic media for 24 hours. The RNA was collected using TRIzol Reagent (Invitrogen, cat. 15596018). Morphology of microglia in acidity tests were analysed from images taken with Zeiss Axio Observer Phase contrast microscope. Cells were annotated manually either into ramified or amoeboid categories. Statistical tests were done using GraphPad Prism (v10.3.1).

### Hypoxia *in vitro* cultures

Cells were cultured in hypoxia chambers in an OxyGenie ™ system (BioGenium Microsystems Oy) with varying conditions for 24 hours. Prior to starting the experiment, cells were allowed to adjust to lower oxygen levels by incubating them overnight in a hypoxia incubator (5% O_2_) before exposing them to lower Oxygen tension (0-5%O_2_). For lower pH-level tests, the cell culture medium was adjusted to desired pH value with 0.1 M HCl after letting the medium equilibrate in a cell culture incubator for 1 hour and adjusted media was then sterile filtered before use.

### RNA sequencing and data analysis

Poly-A enrichment was used to construct mRNA libraries of extracted total RNA from macrophage and microglia *in vitro* cultures. Sequencing was done with NovaSeq X Plus series (PE150) technology, with at least 12Gb of data per sample. For bulk-RNA sequencing data analysis, adapter sequences, and low-quality reads were trimmed with TrimGalore (v0.6.7, https://github.com/FelixKrueger/TrimGalore, accessed on 25 June 2024), and the raw sequence files were quality-controlled with FastQC (v0.11.9) ^55^. STAR (v 2.7.10b) ^56^ was used to align the reads to hg38 and featureCounts function from Rsubread package 2.20.0 ^57^ to form a gene count matrix. ComBat_seq function from sva package (v3.54.0) ^58^ was used to reduce batch effect between experiment rounds. The expression levels were normalized with the DESeq2 (v1.46.0)^59^ in R (v4.4.1). For heatmap visualizations, gene expression data was log2-transformed (log2(counts+1)), and the mean expression was calculated for each hypoxia-acidity group. Each gene was then centered by subtracting its mean expression across all groups. Gene set activity analysis was performed as in ^60^ by first z-scoring gene expression values across samples. The activity score was calculated as the sum of z-scores divided by the square root of the number of genes in the set (genes with zero variance were excluded).

### Oxygen gradient

MG with either neutral or acidic pH and MDMs in neutral pH with high and low cell confluencies were exposed to an oxygen gradient created with a gradient lock system (BioGenium Microsystems Oy). Cells were cultured in the gradient for 24 hours and live imaged with Leica DMi8 at 5-minute imaging intervals to track the cell count and movement.

Cell detection and tracking were performed using TrackMate in Fiji (version 2.14.0) [TRACKMATE][FIJI]. Cells were detected using a Difference of Gaussians detector with a threshold of 300 and an estimated radius of 8.5. All other parameters were kept at their default values. Cells were tracked with the Linear Assignment Problem tracker using default settings. Results were quantified in Python by dividing gradient horizontally into 10 segments and change in cell count during 24 hours in these areas was visualized with GraphPad Prism (v10.3.1).

### Visualizations

Visualizations in R were created using ggplot2 (v4.0.1), ggbreak (v0.1.6), ComplexHeatmap (v2.22.0), RColorBrewer (v1.1-3), ggVennDiagram (v1.5.4), and gplots (v3.2.0), unless otherwise specified.

### Statistical analysis

Statistical testing was performed using the Wilcoxon rank-sum test with Benjamini-Hochberg correction for multiple testing in R (v.4.4.1), unless otherwise mentioned in the methods. Effect sizes were estimated using Cliff’s Delta (rcompanion v2.5.1). The experiments were not randomized, and the investigators were not blinded to allocation while doing the experiment and interpreting the results. No data, except those cIHC hypoxic areas that contained <20 cells, was excluded from the analysis.

## Acknowledgments

We would like to acknowledge Ms. Meeri Pekkarinen and Ms. Anja Hartewig for the preprocessing of DNA methylation data (from Capper et al.). We would like to acknowledge Ms. Päivi Martikainen, Ms. Maria Annala, Mrs. Hanna Selin, Mrs. Sari Toivola for sample handling and logistics. We would like to thank everyone who donated their blood sample for the study and thank Mrs. Kristiina Lehtinen, and Mrs. Janette Hinkka, Ms. Rita-Maria Lehmonen and Ms. Marika Vähä-Jaakkola for assistance in sample collection. We are very grateful to Dr. Teijo Pellinen from the University of Helsinki for sharing his cell neighborhood analysis pipeline and insights related to that. Personnel at Tampere University Hospital and Fimlab Laboratories Ltd. are acknowledged for their contribution to sample collection. We are grateful to them and the patients for permitting the analysis of precious patient material. We are grateful for Histology and Genomics core services of the Faculty of Medicine and Health technology, Tampere University.

## Authors’ contributions

H.H., A.J., K.J.R., and A.M.T. conceptualized the study. K.J.R., A.J. P.K., P.R., S.H., and M.N. supervised this work. A.M.T., T.H., J.T., J.M.K., G.K., S.M., and M.H., performed experimental studies. J.K., M. Marttinen, and K.J., contributed to data analysis tools. I.S., I.K., S.J., M. Mohammadlou, K.J., and M.V. performed computational analyses. H.H., J.B., R.R. provided patient sample material. A.M.T, I.S. and K.J.R. wrote the original manuscript draft. A.M.T. designed the figure panels. A.M.T., I.S., I.K., A.S.R., S.J., T.H., J.M.K., G.K., S.M., M.V., J.H., M.N., S.P., P.K., S.H., J.K., V.M.R., and K.J.R. reviewed and edited the manuscript. All authors reviewed and approved the final version of the manuscript.

## Ethics declarations

The study was conducted in accordance with the principles of the Declaration of Helsinki. This study was approved by The Regional Ethics Committee of Tampere University Hospital (Tays)(decision R07042, renewed 12/2024) and Valvira (V/78697/2017). Supportive statement from The Regional Ethics Committee of Tampere University Hospital received also for the production of the human iPSC line (R08070) and its use in neuronal research (R20159). The hiPSCs were acquired from a voluntary subject who had given written and informed content.

## Data availability

The generated bulk RNA-sequencing data will be made available in GEO upon publishing. GeoMx DSP counts are available in the Supplementary Table 4, and raw data is available with request.

## Code availability

All generated scripts are available at https://github.com/CRI-group/Hypoxia_immunology.

## Materials & Correspondence

Correspondence and material requests should be addressed to the corresponding author Kirsi Rautajoki.

---

**Supplementary Figure 1.**
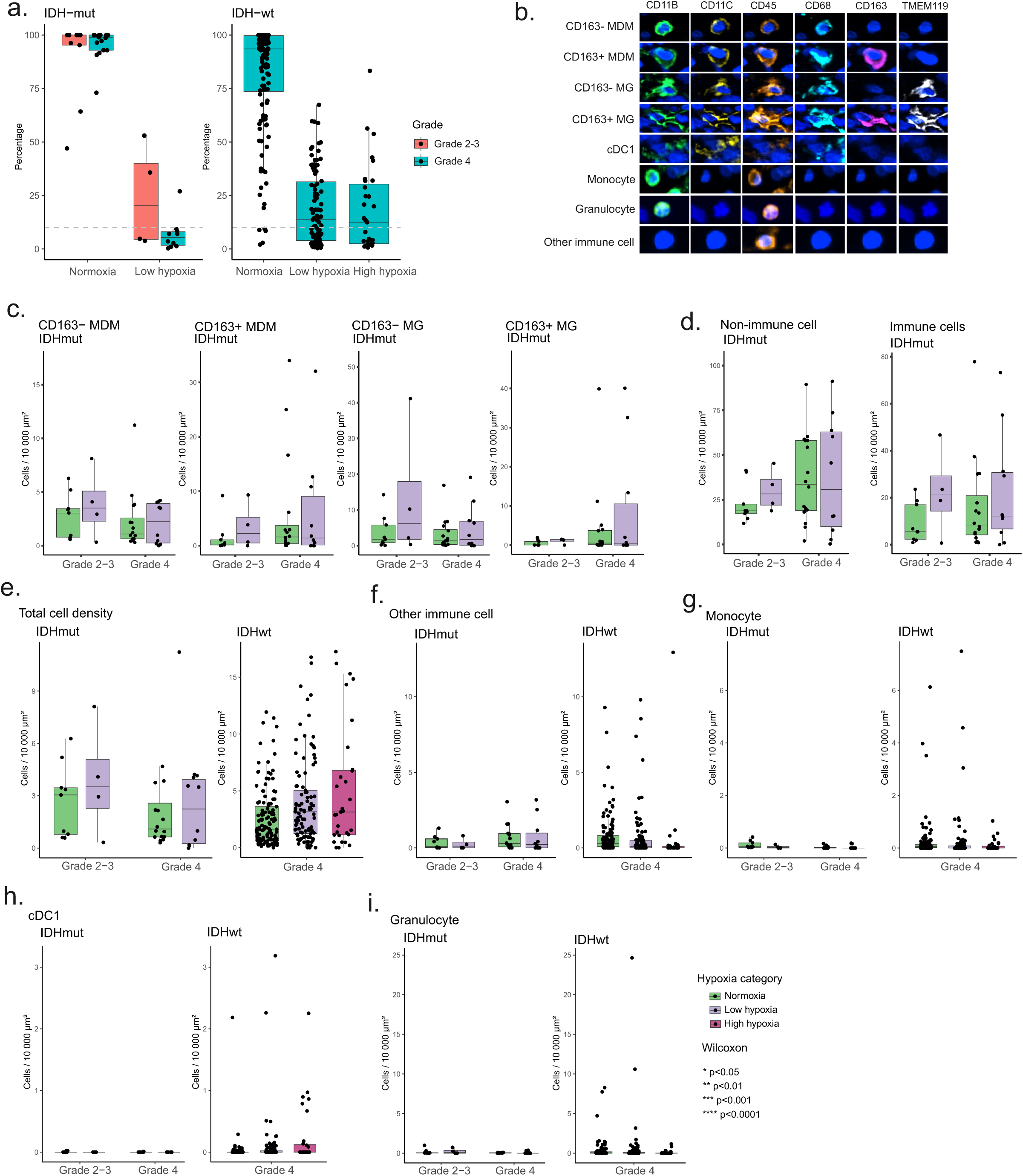
a) Sample-wise pixel fractions across hypoxia categories in both IDHmut astrocytomas and IDHwt GBMs in cIHC dataset. Highly hypoxic regions with over 20 cells were found only in GBMs. b) Representative images of marker intensities and typical cell shapes for each cell type identified from cIHC data with the QuPath cell classifier. c-d) No statistically significant differences were seen in MDM and MG subtypes (c) or in non-immune cell and immune cell densities (d) in IHDmut tumors across hypoxia categories. e) Total cell densities in GBMs and IDHmut astrocytomas across hypoxia categories show no statistically significant differences indicating that hypoxic areas are viable tumor tissue with no distinct drop in cell density. f-i) No statistically significant differences in the densities of Other immune cells (f), Monocytes (g), cDC1s (h), or Granulocytes (i) were detected between different hypoxia categories in either IDHwt GBMs or IDHmut astrocytomas. The Wilcoxon test was used for the statistics in subpanels c-i.

**Supplementary Figure 2.**
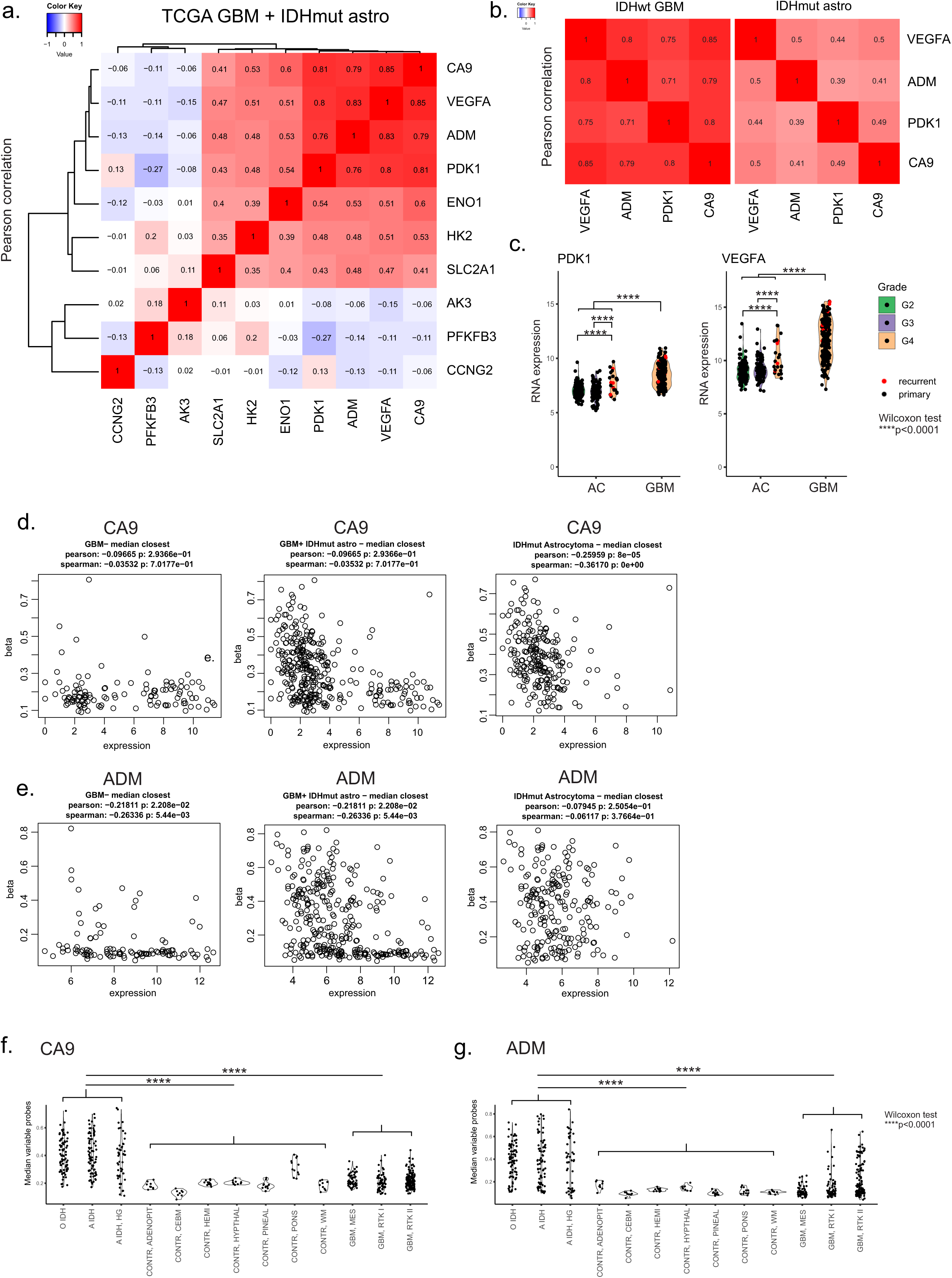
a) Pearson correlation of hypoxia response gene set reported by ^32^ in TCGA GBM and IDHmut astrocytoma (AC) tumors. CA9, ADM, VEGFA and PDK1 show the highest correlation in this dataset. b) Pearson correlation of CA9, ADM, VEGFA and PDK1 in TCGA GBM and IDHmut AC tumors separately. c) Expression of PDK1 and VEGFA in the TCGA GBM and IDHmut AC tumors show significantly higher expression in GBMs compared to ACs (Wilcoxon test, ****p<0,0001). d) Correlation of DNA methylation and RNA expression of CA9 in the TCGA dataset in GBMs, GBMs + IDHmut ACs and in IDHmut ACs separately show negative trends in correlation. e) Correlation of DNA methylation and RNA expression of ADM in the TCGA dataset in GBMs + IDHmut ACs and in IDHmut ACs separately show negative trends in correlation. f) DNA methylation of CA9 in ^33^ dataset shows statistically significant differences in DNA methylation in IDHmut tumors compared to control regions in the brain and GBMs. Similar trend is seen for f) ADM (Wilcoxon test, ****p<0,0001).

**Supplementary Figure 3.**
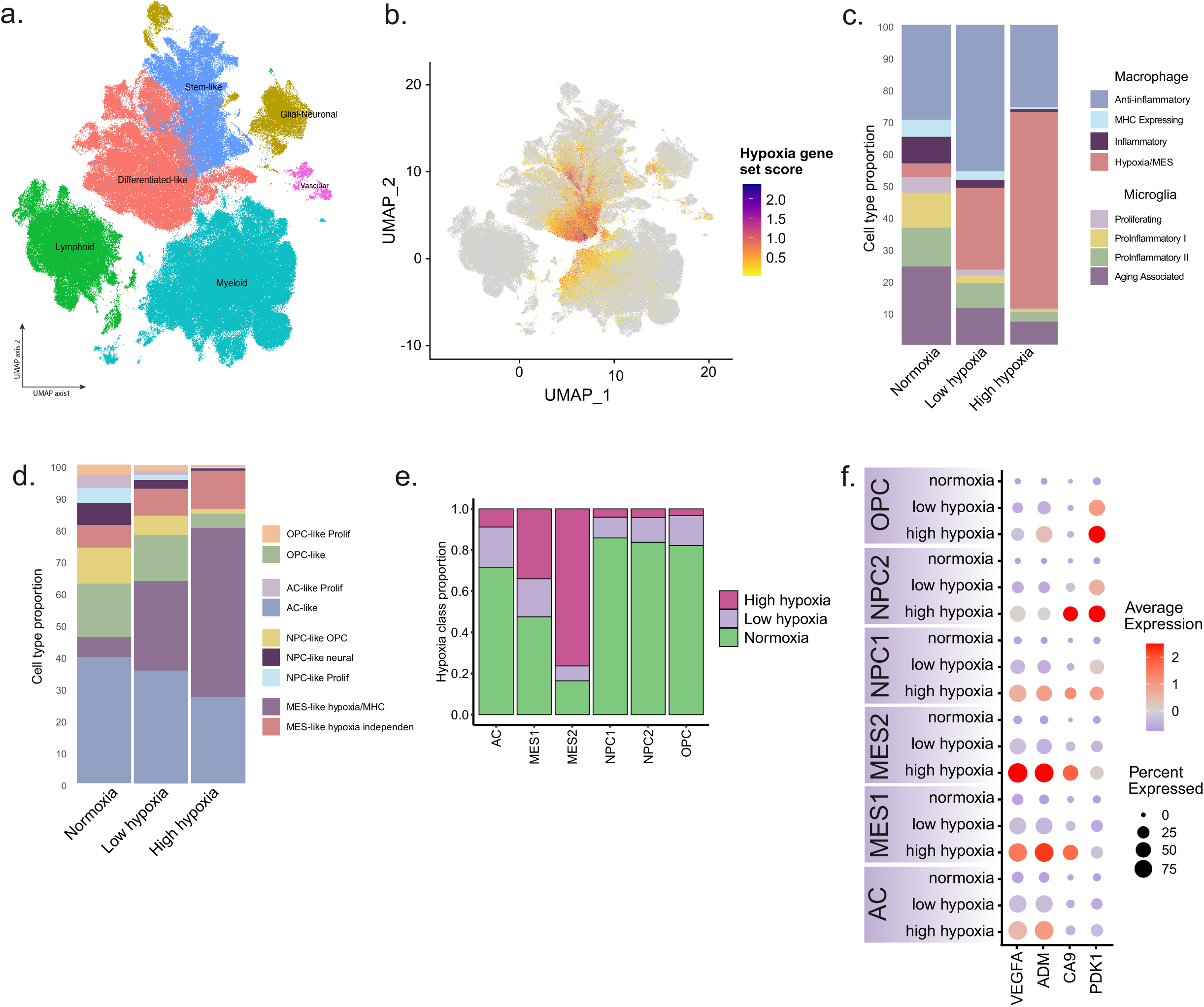
a) Identified major cell types in the GBmap dataset. b) Hypoxia gene set score in the GBmap dataset shows activation mostly in the malignant cell clusters (differentiated and stem-like cells) and myeloid cells. c) Cell type composition of GBmap dataset in hypoxia categories. Hypoxia/MES-subtype of macrophages is especially increased in low and high hypoxia categories whereas MG subsets are depleted. d) A stacked bar plot visualization of cancer cell subtypes across hypoxia categories in GBmap dataset show a major increase of MES-like hypoxia dependent GBM subtype in high hypoxia category. e) A stacked bar plot visualization of cancer cell subtypes across hypoxia categories in our SCS dataset we have assembled from four publications show a major increase of MES-2 subtype in high hypoxia category. f) Hypoxia response genes are generally upregulated under hypoxia in cancer cell subtypes in our SCS dataset we have assembled from four publications. However, cancer cell subtype-associated differences were also detected in the expression patterns.

**Supplementary Figure 4.**
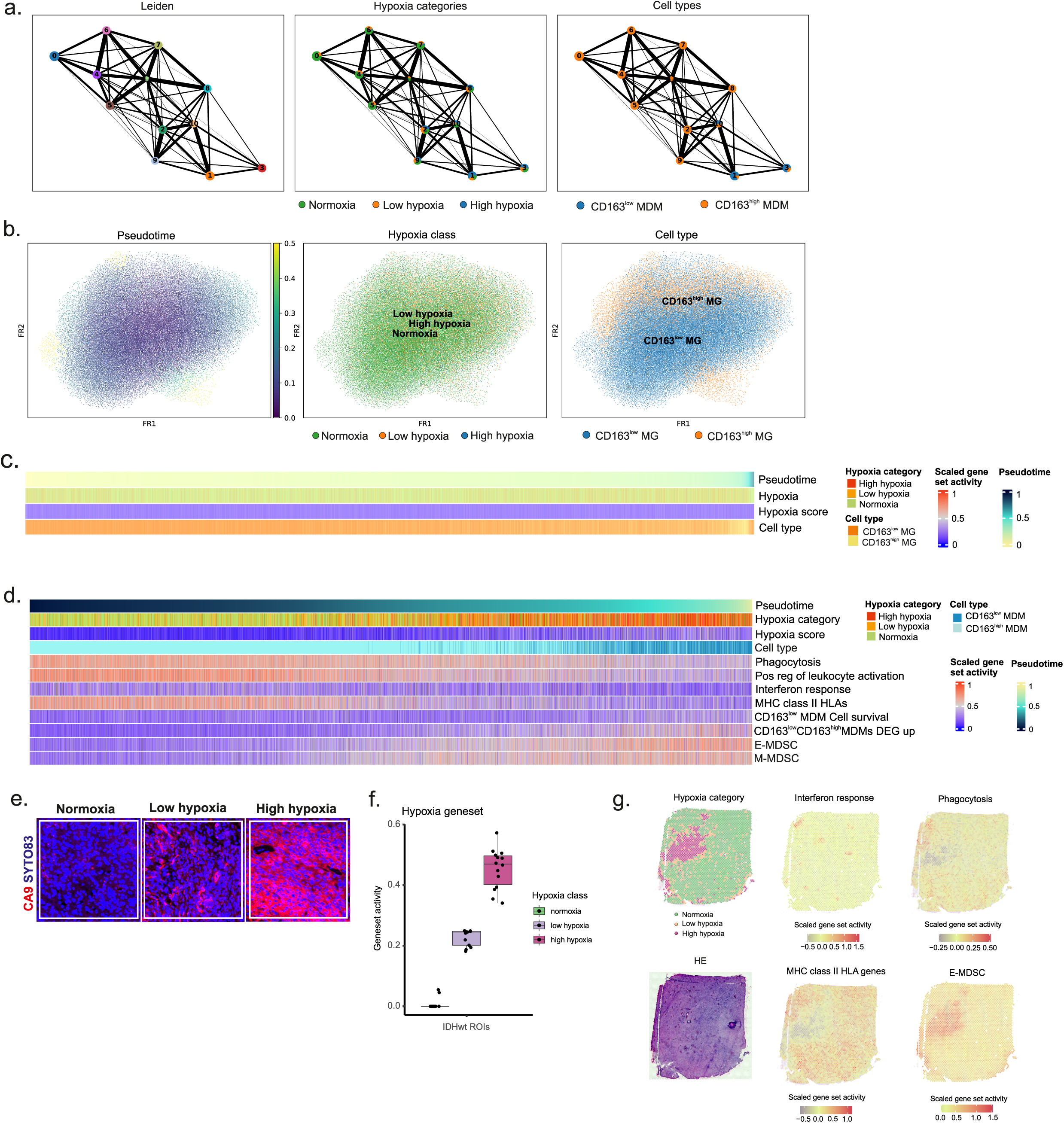
a) Leiden clusters of PAGA done with MDMs using our SCS dataset. Distribution of cells in each cluster to hypoxia categories and subtypes are also shown. b) PAGA pseudotime analysis with only MG shows that hypoxia response in MG subtypes are scattered and not organized similarly as MDMs. c) Hypoxia gene set activity and hypoxia score in pseudotime trajectory of MG shows that increase of hypoxia doesn’t result in shift in MG phenotype, so the trajectory is driven by other factors. d) Gene set activities in PAGA pseudotime trajectory of MDMs without B-spline smoothing. e) Examples of CA9 staining which was used to select ROIs for analysis with GeoMx from the GBM cohort. f) Hypoxia gene set activity was used to categorize GeoMx ROIs into hypoxia categories. g) Representative images of the Activity of MHC class II, Interferon response, Phagocytosis, and CD163^low^ CD163^high^ MDM DEG up gene sets in a GBM sample analyzed with Visium.

**Supplementary Figure 5.**
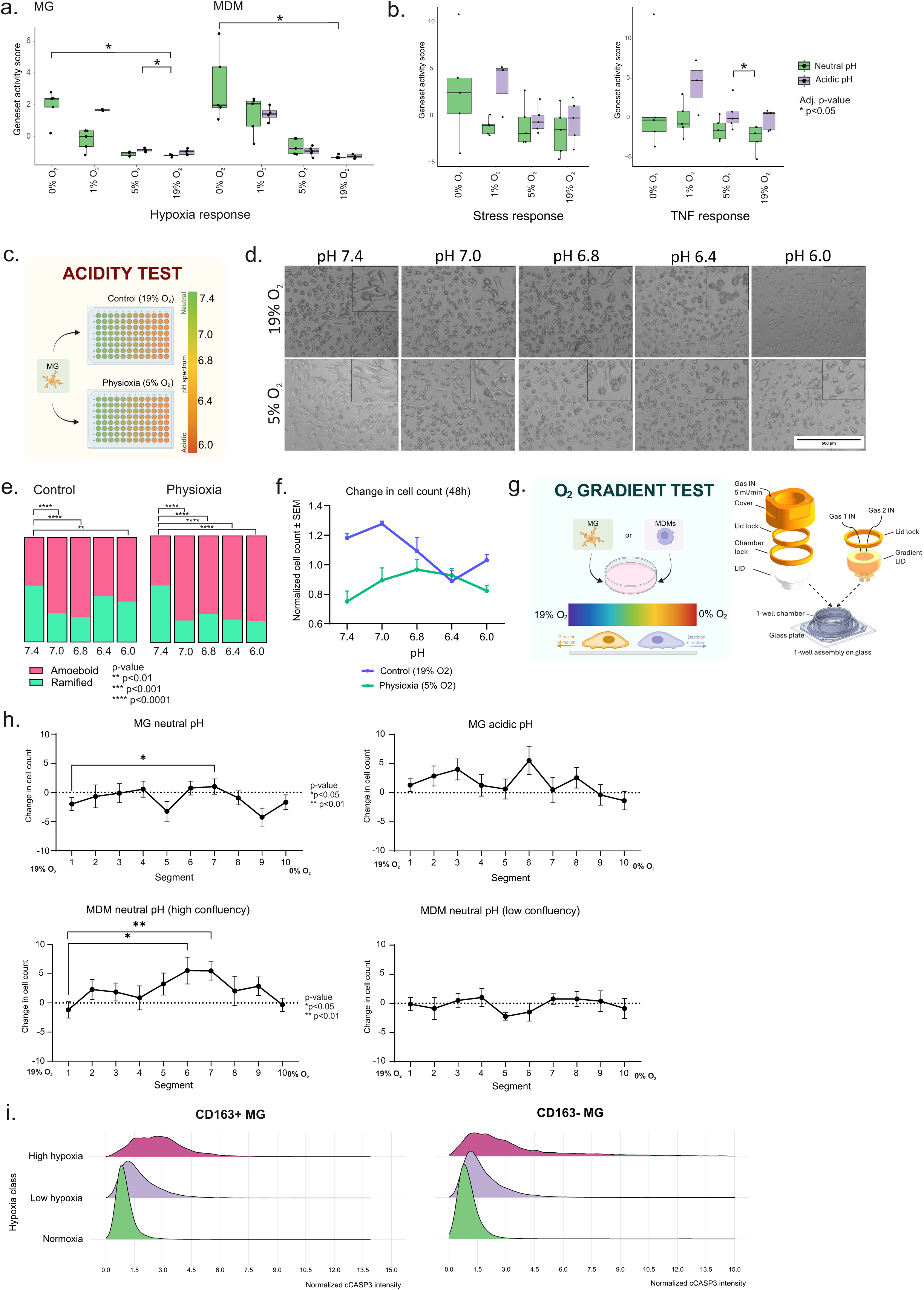
a) Activity of hypoxia response gene sets in MG and MDM *in vitro*. In MG, acidity alone already upregulates the hypoxia response whereas in MDMs upregulation is seen especially in hypoxia (1% O_2_) and anoxia (0% O_2_) and pH level does not seem to affect the level of gene expression (Wilcoxon test). b) Activity of TNF and Stress response gene sets in MG *in vitro*. Acidity induces upregulation of both responses, although statistically significance was not reached. c) Concept figure of acidity test done with MG *in vitro*. MG were exposed to decreasing pH levels in control (19% O_2_) and physioxic (5% O2) oxygen conditions for 48 hours. d) Change in MG cell morphology can be seen from phase contrast microscopy images captured at the end of the acidity test. e) Proportion of MG cell shapes quantified from microscopy images of the acidity test show significant change towards amoeboid shape in acidity, especially when combined with physioxia (Fisher’s exact test). f) Relative change in cell count in MG in the acidity test conditions. Dramatic drop in the cell count in respect to 0 hours was not observed in physioxic and/or acidic conditions. g) Concept figure of gradient test done with both MG and MDM created with Biorender ^34^. Cells were exposed to an oxygen gradient for 24 hours. MG were exposed with either neutral or acidic pH and MDMs with neutral pH with either high or low cell confluency. Schematic illustration of equipment that were used to create hypoxic/anoxic and oxygen gradients for the cells in Hypoxia and Gradient tests. h) For either MDMs nor MG, the change in the oxygen level did not result in significant change in cell confluence along the gradient, although the wide oxygen range can contribute to the results. Oxygen gradient results were quantified by dividing the gradient area horizontally into 10 segments (from 0% to 19%, so with approximately 2 % intervals) to identify distinct changes in cell migration or accumulation. Also, no preference in the direction of cell migration was observed with visual inspection. i) Normoxia-normalized cCASP3 intensity grows in both microglia subtypes as level of hypoxia increases.

